# Evaluating the Immunological cross-reactivity of Indian polyvalent antivenoms towards the venom of *Hypnale hypnale* (hump-nosed pit viper) from the Western Ghats

**DOI:** 10.1101/2020.08.01.232579

**Authors:** Muralidharan Vanuopadath, Dileepkumar Raveendran, Bipin Gopalakrishnan Nair, Sudarslal Sadasivan Nair

## Abstract

*Hypnale hypnale* (hump-nosed pit viper) is a venomous pit viper species found in the Western Ghats of India and Sri Lanka. Due to the severe life-threatening envenomation effects induced by its venom components, *Hypnale hypnale* has been classified under ‘category 1’ of medically important snake species by the World Health Organization. Since there are no specific antivenoms available to combat its envenomation in India, the only option available is to administer Indian polyvalent antivenoms. However, the cross-neutralization potential of the commercially available polyvalent antivenoms on Indian *Hypnale hypnale* venom has not been explored so far. In the current study, in vitro immunological cross-reactivity of *Hypnale hypnale* venom towards various Indian polyvalent antivenoms were assessed using end point titration ELISA and Western blotting. A three to four-fold increase in EC_50_ values were obtained for *Hypnale hypnale* venom towards all the antivenoms tested. Observation of minimal binding specificities towards low and high molecular mass venom proteins are suggestive of the fact that commercially available polyvalent antivenoms failed to detect and bind to the antigenic epitopes of considerable number of proteins present in *Hypnale hypnale* venom. This highlights the importance of including *Hypnale hypnale* venom in the immunization mixture while raising antivenoms.

## Introduction

Snakebite has been included in the neglected tropical disease category by the World Health Organization (WHO) in 2017 because of the associated mortality and morbidity cases [1]. A nation-wide mortality survey estimated that approximately 1.2 million snakebite deaths occurred from 2000 to 2019 accounting for 58,000 deaths/year in India [2]. The only treatment option available against envenomation is to administer the desired amount of antivenoms at the right time. Delay in providing antivenoms, misidentification of snake species and improper treatment modalities increase the mortality rates. Despite these, lack of appropriate antivenoms also accounts for most of the mortality or morbidity cases [1]. In India, antivenoms are raised against the ‘big four’ snake species, *Daboia russelii* (Russell’s viper), *Echis carinatus* (saw-scaled viper), *Bungarus caeruleus* (common krait) and *Naja naja* (spectacled cobra). Since these four snake species are widely distributed across the country, majority of the snakebites and envenomation cases reported are due to their bites [3, 4]. Apart from these ‘big four’, India is abode to many other medically-relevant snakes including several species of kraits, cobras, pit vipers and sea snakes [5]. However, to date, there are no specific antivenoms available towards their envenomation.

In India, the antivenom manufacturers solely depend on the Irula Snake Catchers Industrial Cooperative Society (Tamilnadu, South-India) for obtaining venoms used for the immunization process [4]. However, several in vitro preclinical studies indicate that the immuno-recognition potential of these antivenoms towards the venom toxins present even in the ‘big four’ snakes from different geographical locations seems to be varied [6-9]. Besides these, studies have shown that available Indian polyvalent antivenoms are not effective in neutralizing venom components present in the ‘non-big four’, yet medically-relevant snake species including *Trimeresurus malabaricus, Naja kaouthia, Echis carinatus sochureki, Bungarus fasciatus* and *Bungarus sindanus* [10, 11].

*Hypnale hypnale* (hump-nosed pit viper; HPV) belongs to the ‘non-big four’ group and is distributed across Sri Lanka and the Western Ghats of India [5]. Due to the severity of complications associated with its bite, HPV has been classified under the ‘category 1’ of medically important snake species by WHO [3]. Clinical reports from Kerala (a state in the southern part of India) suggest that one-fourth of the viper bites are caused due to HPV and are often misidentified to that of saw-scaled viper bites [3, 12, 13]. However, apart from the polyvalent antivenoms raised against ‘big four’, currently there are no specific antivenoms available against its envenomation. Interestingly, none of the available reports demonstrate the immunological cross-reactivity profiles of Indian polyvalent antivenoms against HPV venom. In the current study, through endpoint titration ELISA and western blotting, our main objective is to determine the immunological cross-reactivity profiles of the major Indian polyvalent antivenoms against the venom of HPV from the Western Ghats.

## Materials and methods

Methanol, acrylamide, bisacrylamide, ammonium persulphate, coomassie brilliant blue R-250 (CBB R-250), beta-mercaptoethanol, bromophenol blue, glycerol, glycine, Tetramethyl benzidine/Hydrogen Peroxide (TMB/H_2_O_2),_ HRP-conjugated secondary antibody and Tetramethylethylenediamine were purchased from Sigma-Aldrich. ECL substrate, Tris, Sodium dodecyl sulfate (SDS) and PVDF membrane were procured from Bio-Rad. The protein ladder was purchased from Thermo Fisher Scientific. All other chemicals and reagents used were either of analytical or LC-MS grade.

### Venom collection

*Hypnale hypnale* and *Echis carinatus* (Saw-scaled viper; SSV) venoms from the Western Ghats of India were collected and stored as described earlier [14].

### Antivenoms

Lyophilized equine antivenoms raised against the ‘big four’ snakes of India (*Echis carinatus, Naja naja, Bungarus caeruleus*, and *Daboia russelli*) obtained from VINS (batch number: 01AS16040, expiry date: 07/2020), Virchow (batch number: PAS00116, expiry date: 01/2020) and Premium Serums and Vaccines (PSAV; batch number: ASVS (I) Ly-010, expiry date: 08/2022) were used for the entire immunological cross-reactivity assays.

### End point titration ELISA

For performing ELISA, 1000 ng of HPV and SSV venoms were added on to a high bind ELISA plate (Costar) and kept for overnight incubation at 4°C. The unbound venoms were removed from the wells by inverting the plates and flick drying. Followed by this, any non-reactive sites in the wells were blocked by adding 2.5% BSA in phosphate-buffered saline (PBS) and incubated for 2 h at room temperature (RT). The primary antibodies were serially diluted using 2.5% BSA by a factor of three (1:100, 1:300, 1:900, 1:2700, 1:8100, 1:24300, 1:72900, 1:218700, 1:656100, 1:1968300, 1: 5004900) and was incubated for 2 h at RT. The unbound antibodies were removed through 0.1% tween-20 in phosphate-buffered saline (PBST) wash and this step was repeated thrice. Followed by this, secondary antibody (HRP conjugated) at a dilution of 1:32,000 in 2.5 % BSA was added on to each well and kept for incubation at RT for 2 h. Excess of secondary antibody was removed through 0.1% PBST wash as mentioned above. To the wells, 100 µL of HRP substrate solution, TMB/H_2_O_2_, was added and incubated in the dark for 30 min at RT. The endpoint readings were taken by terminating the reaction by adding 100 µL of 0.5 M sulphuric acid to each well. The optical density values were monitored at 450 nm using a plate reader. The EC_50_ values were computed from the obtained OD values through non-linear regression analysis using Prism (version 6.01) software.

### Western blotting

For immunoblotting experiments, 50 µg of crude HPV and SSV venom components resolved on 15% SDS gel were transferred (25 V for 1 h) to a PVDF membrane. To minimize non-specific binding, the membrane was blocked using 5% BSA at RT for 1 h. Antivenoms obtained from Virchow, PSAV and VINS were used as primary antibodies (dilution used, 1:500) and was kept for incubation at 4 °C (overnight). The membrane was washed thrice for 10 min using 0.1% PBST solution. To this, secondary antibody (HRP conjugated) at 1:5000 dilution in 5% BSA was added and kept at RT for 1 h. The excess of secondary antibody was then removed through 0.1% PBST wash as mentioned above. Followed by this, the ECL substrate was added on to the membrane and signals indicating the venom-antivenom interaction were captured using a gel documentation system (Bio-Rad).

### Statistical analysis

Prism (GraphPad Software Inc., San Diego, CA; version 6.01) was used for performing statistical comparisons. The data obtained from triplicate experiments were plotted with the help of the software and are represented as mean ± standard deviation or as stated, otherwise.

## Results and discussion

*Hypnale hypnale* shares striking physical resemblances to that of the saw-scaled viper except the scaling patterns seen on their heads. Besides, the clinical complications associated after HPV bites match that of saw-scaled vipers’ [13, 15]. These similarities prompted us to compare the immuno-reactivity patterns of HPV and SSV venoms against the Indian polyvalent antivenoms using SSV venom as a positive control. The end point titration ELISA performed using the three antivenoms suggest that the antigenic epitopes present in HPV venom were recognized less efficiently than that in the SSV venom (Figure 1a-c). It is evident that, since SSV venom was included in the immunization mixture while raising the antivenoms [3], the protein components present in SSV venom showed considerable immunological cross-reactivity profiles towards all the antivenoms. As seen in Figure 1 (a-c; insets), compared to SSV venom, all the antivenoms tested against HPV venom were showing a 3-4-fold increase in the EC_50_ values, (18.23, 21.61 and 21.79 µg/mL for VINS, PSAV and Virchow, respectively) suggesting that it would require more amount of antivenoms in neutralizing the toxin components present. The higher EC_50_ values obtained for HPV venom against all the tested antivenoms corroborate the clinical findings indicating the inefficiency of antivenoms in treating HPV envenomed victims and any attempts to administer more amount of antivenoms result in adverse reactions than positive outcomes [13, 15, 16]. Therefore, a general strategy the clinicians follow for HPV envenomation is subjecting the victims to continuous dialysis and ventilation instead of antivenom administration [13, 15]. Further, as seen in Figure 1, the end point dilution values for VINS and Virchow antivenoms against HPV venom were 1:24,300 and that of PSAV was 1:8,100. But the minimum dilution factor required for all the tested antivenoms in detecting the SSV venom components was same at 1:72,900. The data also revealed that VINS and Virchow antivenoms evoke comparatively better immuno-reactive profiles against HPV venom than PSAV antivenom. This suggests that the antivenom immunological cross-reactivity profiles vary within vendors also. The production and purification strategies followed by the vendors during antivenom formulation might be a critical factor that influences the cross-reactivity [16].

**Figure 1.**
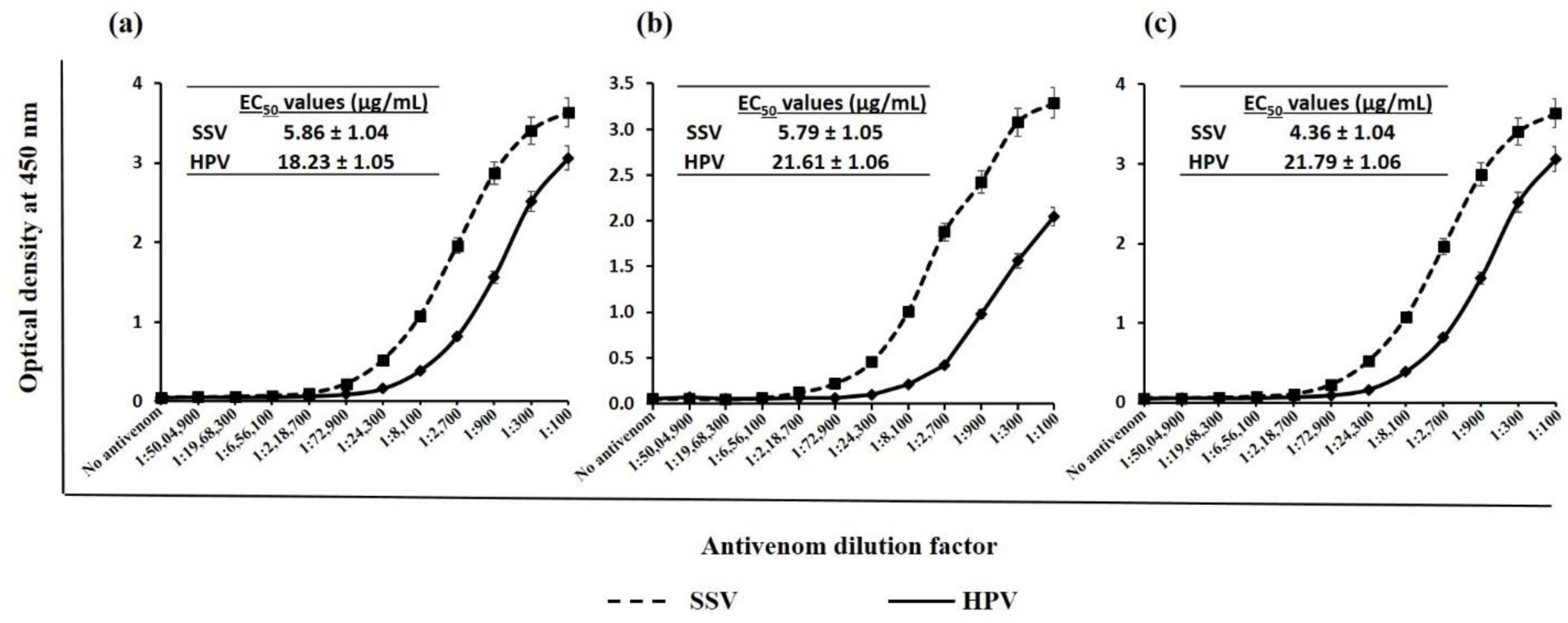
Immunological cross-reactivity of SSV and HPV venoms assessed through end point titration ELISA. Titration curves obtained using various dilutions of (a) VINS (b) PSAV and (c) Virchow antivenoms. The computed EC_50_ values are shown in the insets. (SSV: saw-scaled viper (positive control); HPV: hump-nosed pit viper)

Subsequent to ELISA, immunoblot analysis were performed to compare the binding specificities of SSV and HPV venom components against the major Indian antivenom preparations. Though SDS-PAGE analysis (Figure 2a) reveal presence of low and high molecular mass proteins in SSV and HPV venoms, the western-blotting indicate that all the tested antivenoms bind less efficiently to low and high molecular mass proteins present in HPV venom than that in the SSV (Figure 2 b-d). The inadequacy of polyvalent antivenoms in detecting low and high molecular mass proteins present in different snake species has been reported by various groups, which is in agreement with our finding [6-10]. Further, we have observed that the binding specificity of VINS and PSAV antivenoms (Figure 2b & c) were better than Virchow antivenom (Figure 2d) in detecting HPV venom epitopes. Similar trend was noticed in detecting SSV venom epitopes also. As stated above, the differences in the antivenom neutralization efficiencies against HPV and SSV venom might be contributed by the antivenom production and purification protocols and the variation in the venom antigens that were used for immunizing horses while generating polyvalent antivenoms [16, 17]. In short, our results support the need to include HPV venom also in the immunization mixture during the production of antivenom in India. On a similar note, a pre-clinical study from Sri Lanka on a new poly-specific antivenom has shown to have improved neutralization efficiency against Sri Lankan HPV venom when it was part of the immunization mixture [18].

**Figure 2.**
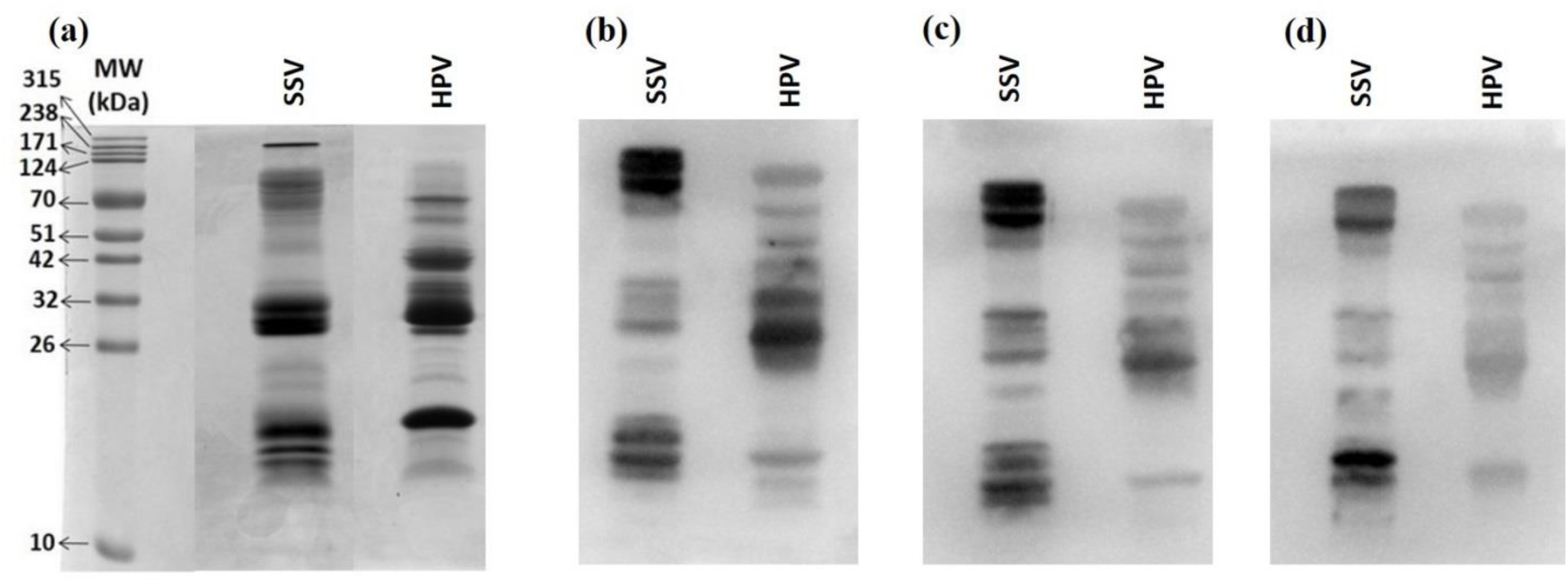
Immunological cross-reactivity of SSV and HPV assessed through Western blotting. (a) HPV and SSV venoms resolved on a 15% SDS-PAGE gel under reducing conditions. Western blotting of the venoms using (b) VINS (c) PSAV and (d) Virchow polyvalent antivenoms. (SSV: saw-scaled viper; HPV: hump-nosed pit viper)

## Acknowledgments

The authors would like to acknowledge Sri. Mata Amritanandamayi Devi, The Chancellor of Amrita Vishwa Vidyapeetham, for being the inspiration behind this study. We are grateful to the forest department of Kerala for their assistance during the field visits during the time of venom collection. MV thank Amrita Vishwa Vidyapeetham and Amrita Centre for Research and Development for providing funding in the form of fellowship. Financial assistance in executing this project provided by Amrita Vishwa Vidyapeetham and Indriyam Biologics is also duly acknowledged. The infrastructural support provided by Amrita Agilent Analytical Research Centre is also duly acknowledged.

## References

[1] J.P. Chippaux: Snakebite envenomation turns again into a neglected tropical disease! J Venom Anim Toxins Incl Trop Dis 2017, 23: 38.doi:10.1186/s40409-017-0127-6

[2] W. Suraweera, D. Warrell, R. Whitaker, G. Menon, R. Rodrigues, S.H. Fu, R. Begum, P. Sati, K. Piyasena, M. Bhatia, P. Brown, P. Jha: Trends in snakebite deaths in India from 2000 to 2019 in a nationally representative mortality study. Elife 2020, 9: 10.7554/eLife.54076

[3] I.D. Simpson, R.L. Norris: Snakes of medical importance in India: is the concept of the “Big 4” still relevant and useful? Wilderness Environ Med 2007, 18: 2–9.doi:10.1580/06-weme-co-023r1.1

[4] D.A. Warrell, J.M. Gutierrez, J.J. Calvete, D. Williams: New approaches & technologies of venomics to meet the challenge of human envenoming by snakebites in India. Indian J Med Res 2013, 138: 38–59.

[5] R. Whitaker, G. Martin: Diversity and Distribution of Medically Important Snakes of India. In: Gopalakrishnakone P, Faiz A, Fernando R, Gnanathasan C A, Habib A G, Yang C-C, editors. Clinical Toxinology in Asia Pacific and Africa. Toxinology; Springer, Dordrecht 2015, 2: 115–136.doi: https://doi.org/10.1007/978-94-007-6386-9_16

[6] A. Patra, A. Chanda, A.K. Mukherjee: Quantitative proteomic analysis of venom from Southern India common krait (Bungarus caeruleus) and identification of poorly immunogenic toxins by immune-profiling against commercial antivenom. Expert Rev Proteomics 2019, 16: 457–469.doi:10.1080/14789450.2019.1609945

[7] B. Kalita, A. Patra, A.K. Mukherjee: Unraveling the Proteome Composition and Immuno-profiling of Western India Russell’s Viper Venom for In-Depth Understanding of Its Pharmacological Properties, Clinical Manifestations, and Effective Antivenom Treatment. J Proteome Res 2017, 16: 583–598.doi:10.1021/acs.jproteome.6b00693

[8] A. Patra, B. Kalita, A. Chanda, A.K. Mukherjee: Proteomics and antivenomics of Echis carinatus carinatus venom: Correlation with pharmacological properties and pathophysiology of envenomation. Sci Rep 2017, 7: 17119.doi:10.1038/s41598-017-17227-y

[9] A. Chanda, B. Kalita, A. Patra, W. Senevirathne, A.K. Mukherjee: Proteomic analysis and antivenomics study of Western India Naja naja venom: correlation between venom composition and clinical manifestations of cobra bite in this region. Expert Rev Proteomics 2019, 16: 171–184.doi:10.1080/14789450.2019.1559735

[10] M. Vanuopadath, S.K. Shaji, D. Raveendran, B.G. Nair, S.S. Nair: Delineating the venom toxin arsenal of Malabar pit viper (Trimeresurus malabaricus) from the Western Ghats of India and evaluating its immunological cross-reactivity and in vitro cytotoxicity. Int J Biol Macromol 2020, 148: 1029–1045.doi:10.1016/j.ijbiomac.2020.01.226

[11] R.R. Senji Laxme, S. Khochare, H.F. de Souza, B. Ahuja, V. Suranse, G. Martin, R. Whitaker, K. Sunagar: Beyond the ‘big four’: Venom profiling of the medically important yet neglected Indian snakes reveals disturbing antivenom deficiencies. PLoS Negl Trop Dis 2019, 13: e0007899.doi:10.1371/journal.pntd.0007899

[12] K.S. Kumar, S. Narayanan, V. Udayabhaskaran, N.K. Thulaseedharan: Clinical and epidemiologic profile and predictors of outcome of poisonous snake bites - an analysis of 1,500 cases from a tertiary care center in Malabar, North Kerala, India. Int J Gen Med 2018, 11: 209–216.doi:10.2147/IJGM.S136153

[13] J.K. Joseph, I.D. Simpson, N.C. Menon, M.P. Jose, K.J. Kulkarni, G.B. Raghavendra, D.A. Warrell: First authenticated cases of life-threatening envenoming by the hump-nosed pit viper (Hypnale hypnale) in India. Trans R Soc Trop Med Hyg 2007, 101: 85–90.doi:10.1016/j.trstmh.2006.03.008

[14] M. Vanuopadath, N. Sajeev, A.R. Murali, N. Sudish, N. Kangosseri, I.R. Sebastian, N.D. Jain, A. Pal, D. Raveendran, B.G. Nair, S.S. Nair: Mass spectrometry-assisted venom profiling of Hypnale hypnale found in the Western Ghats of India incorporating de novo sequencing approaches. Int J Biol Macromol 2018, 118: 1736–1746.doi:10.1016/j.ijbiomac.2018.07.016

[15] C.A. Ariaratnam, V. Thuraisingam, S.A. Kularatne, M.H. Sheriff, R.D. Theakston, A. de Silva, D.A. Warrell: Frequent and potentially fatal envenoming by hump-nosed pit vipers (Hypnale hypnale and H. nepa) in Sri Lanka: lack of effective antivenom. Trans R Soc Trop Med Hyg 2008, 102: 1120–6.doi:10.1016/j.trstmh.2008.03.023

[16] G. Leon, M. Herrera, A. Segura, M. Villalta, M. Vargas, J.M. Gutierrez: Pathogenic mechanisms underlying adverse reactions induced by intravenous administration of snake antivenoms. Toxicon 2013, 76: 63–76.doi:10.1016/j.toxicon.2013.09.010

[17] R. Teixeira-Araujo, P. Castanheira, L. Brazil-Mas, F. Pontes, M. Leitao de Araujo, M.L. Machado Alves, R.B. Zingali, C. Correa-Netto: Antivenomics as a tool to improve the neutralizing capacity of the crotalic antivenom: a study with crotamine. J Venom Anim Toxins Incl Trop Dis 2017, 23: 28.doi:10.1186/s40409-017-0118-7

[18] M. Villalta, A. Sanchez, M. Herrera, M. Vargas, A. Segura, M. Cerdas, R. Estrada, I. Gawarammana, D.E. Keyler, K. McWhorter, R. Malleappah, A. Alape-Giron, G. Leon, J.M. Gutierrez: Development of a new polyspecific antivenom for snakebite envenoming in Sri Lanka: Analysis of its preclinical efficacy as compared to a currently available antivenom. Toxicon 2016, 122: 152–159.doi:10.1016/j.toxicon.2016.10.007

